# Conformational Ensembles of Non-Coding Elements in the SARS-CoV-2 Genome from Molecular Dynamics Simulations

**DOI:** 10.1101/2020.12.11.421784

**Authors:** Sandro Bottaro, Giovanni Bussi, Kresten Lindorff-Larsen

## Abstract

The 5′ untranslated region (UTR) of the severe acute respiratory syndrome coronavirus 2 (SARS-CoV-2) genome is a conserved, functional and structured genomic region consisting of several RNA stem-loop elements. While the secondary structure of such elements has been determined experimentally, their three-dimensional structures are not known yet. Here, we predict structure and dynamics of five RNA stem loops in the 5′-UTR of SARS-CoV-2 by extensive atomistic molecular dynamics simulations, more than 0.5ms of aggregate simulation time, in combination with enhanced sampling techniques. We compare simulations with available experimental data, describe the resulting conformational ensembles, and identify the presence of specific structural rearrangements in apical and internal loops that may be functionally relevant. Our atomic-detailed structural predictions reveal a rich dynamics in these RNA molecules, could help the experimental characterisation of these systems, and provide putative three-dimensional models for structure-based drug design studies.

## Introduction

The genome of the SARS-CoV-2 virus is a single, positive-stranded RNA consisting of approximately 30 thousand nucleotides. Coding regions of the genome are translated into several non-structural proteins, as well as in the widely studied spike protein, envelope protein, membrane protein, and others [1]. The 5’ and 3’ untranslated regions (UTR) contain cis-acting sequences required for viral transcription and replication [2, 3]. In particular, the 5’-UTR harbours conserved, functional elements that enhance viral transcription [4, 5, 6], and are involved in discontinuous transcription leading to leader-body fusion [7]. Recent studies suggest that the 5’-UTR plays a key role in liquid-liquid phase separation phenomena with the SARS-CoV-2 nucleocapsid protein [8, 9]. Proof-of-principle studies using locked nucleic acids targeting conserved structural motifs in the SARS-CoV-2 genome showed the potential for inhibiting growth via RNA-interacting molecules [10].

Chemical probing experiments, NMR chemical shift measurements, and computational predictions have shown that the 5’-UTR is highly structured [11, 12, 13, 14, 15, 10], and consists of several stem-loop elements (Fig. 1). The secondary structures of the isolated elements are in close agreement with the full-length construct [11], suggesting that they can fold independently. The three-dimensional structure of stem loop 2 (SL2) from SARS-CoV-1, which is completely conserved in SARS-CoV-2, has been determined by NMR spectroscopy [16], while no experimental structures exist for the remaining elements in the 5’-UTR. In a recent computational study [17], Rangan et al predicted the three-dimensional fold of all stem-loop elements in the 5’-UTR, 3’-UTR, and of the frameshifting stimulatory element using Rosetta’s FARFAR2 algorithm [18, 19]. The predicted model of SL2 in the 5’-UTR was later refined using MD simulations, [20] leading to an improved agreement with available NMR data [16].

**Figure 1.**
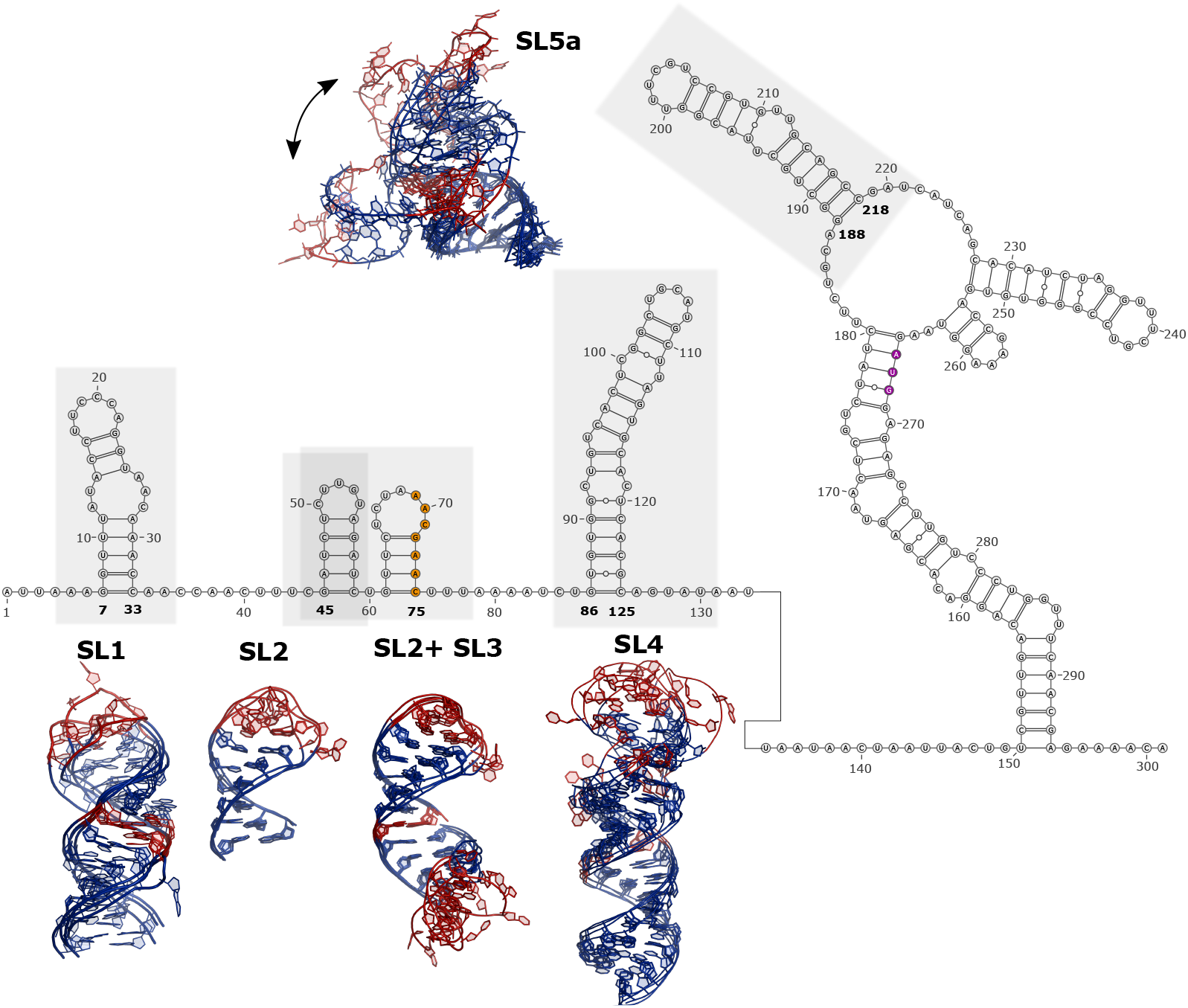
Reference sequence and secondary structure of the 5’-UTR in SARS-CoV-2 genome. The numbering follows the convention in [11]. The five structured elements considered in this work are labeled and highlighted in grey: SL1, SL2, SL2+SL3, SL4, and SL5a. The transcription regulatory sequence and the AUG start codon are highlighted in orange and purple, respectively. Selected centroids of the most populated clusters from MD simulations corresponding to each element are shown.

While there has been a plethora of structural studies of SARS-CoV-2 proteins, much less is known about the structure of the RNA genome at atomic-resolution. RNA molecules often display a rich, complex dynamical behaviour that is difficult to capture using experiments alone and which can play crucial roles in cellular functions [21, 22]. In this work we have thus used atomistic molecular dynamics simulations to predict the structure and dynamics of five 5’-UTR stem-loops. These elements were chosen because their high degree of conservation in betacoronaviruses, together with their vital role in viral replication, make them attractive, potential targets for small molecules. Experimental studies on the same constructs are currently being pursued within the COVID-19 NMR initiative (https://covid19-nmr.de/). The results presented in this work therefore represent a step in using computation and experiments in a synergistic manner. At the same time, the elements that we have chosen to model represent the largest structural elements for which it is currently feasible to obtain relatively converged simulations even with multi-microsecond enhanced sampling simulations.

With this spirit, we have generated three-dimensional configurations compatible with the secondary structure recently determined by chemical probing and NMR experiments [11]. While several fragment-based tools exist to perform such task [23, 24], we here adapt an MD-based procedure [25]. Similar secondary structure restraining approaches have been employed in allatom RNA folding simulations [26, 27] and to study internal loop dynamics [28]. These initial structures are subject to extensive MD sampling, amounting to more than 0.5ms of aggregate simulation time. More precisely, we enhance the sampling of apical and internal loop regions using a temperature replica-exchange scheme in which only specific portions of the RNA solute are affected. [29, 30, 31]. For such regions we currently lack detailed structural information, and available experimental data suggest that conformational rearrangements are more likely to occur. Our simulations are thus a Boltzmann-weighted ensemble of three-dimensional conformations constituting our prediction of structure and conformational heterogeneity of these elements.

First, we apply the computational protocol to SL2, and validate the accuracy of our prediction against the three-dimensional structure of SL2 from SARS-CoV-1 and the NMR data used to generate this structure [16]. We then proceed by describing our predictions on four elements of increasing complexity (Fig. 1), and discuss the agreement with available NMR measurements [32, 33]. Stem loop 5a (SL5a) is part of the four-way junction SL5 containing the AUG starting codon, whose function is linked to RNA packaging [34]. Stem loop 1 (SL1) has been suggested to play a role in escaping the viral mechanism inhibiting mRNA translation [35, 6]. The function of stem loop 4 (SL4) is still not completely understood, and may act as a spacing element [36]. Finally, the SL2+SL3 construct consists of SL2 and stem loop 3 (SL3). The latter element contains the transcription regulatory sequence necessary for discontinuous RNA synthesis [37].

Our predictions could help the atomic-detailed experimental characterisation of such systems, as simulations can be easily integrated a posteriori with NMR or SAXS data [38, 39, 40]. The predicted models provide a starting point for structure-based drug design studies [41, 42], and could serve as additional benchmark systems to evaluate the accuracy of atomistic RNA force-field [43]. To maximize reproducibility and enable others to build on our work [44], we have made trajectories, input files, and analysis scripts available via github at https://github.com/KULL-Centre/papers/tree/master/2020/COVID_5UTR_MD.

## Results and Discussion

In this section we describe the conformational ensembles obtained from MD simulations. We first show the results obtained on SL2, the simplest system considered in this study. Crucially, in this case it is possible to validate the procedure by comparing with available structural data and NMR measurements.

### Stem Loop 2

We run unbiased MD simulations of a single-stranded, extended RNA with sequence GGAUCUCUUGUAGAUCU. After 10ns, we apply a biasing force (in a process we term ‘pulling’) to promote the formation of an ideal A-form RNA structure in the region where the secondary structure is known (blue nucleotides in Fig. 2a). By construction the structural dissimilarity from the template, here evaluated using a nucleic-acids specific distance called ellipsoidal root mean square distance (eRMSD)[45], decreases during pulling (Fig. 2b, top panel). After 40 ns the external bias is removed, and the system fluctuates around a stable structure in which all five Watson-Crick (WC) base-pairs are formed. (Fig. 2b, bottom panel).

**Figure 2.**
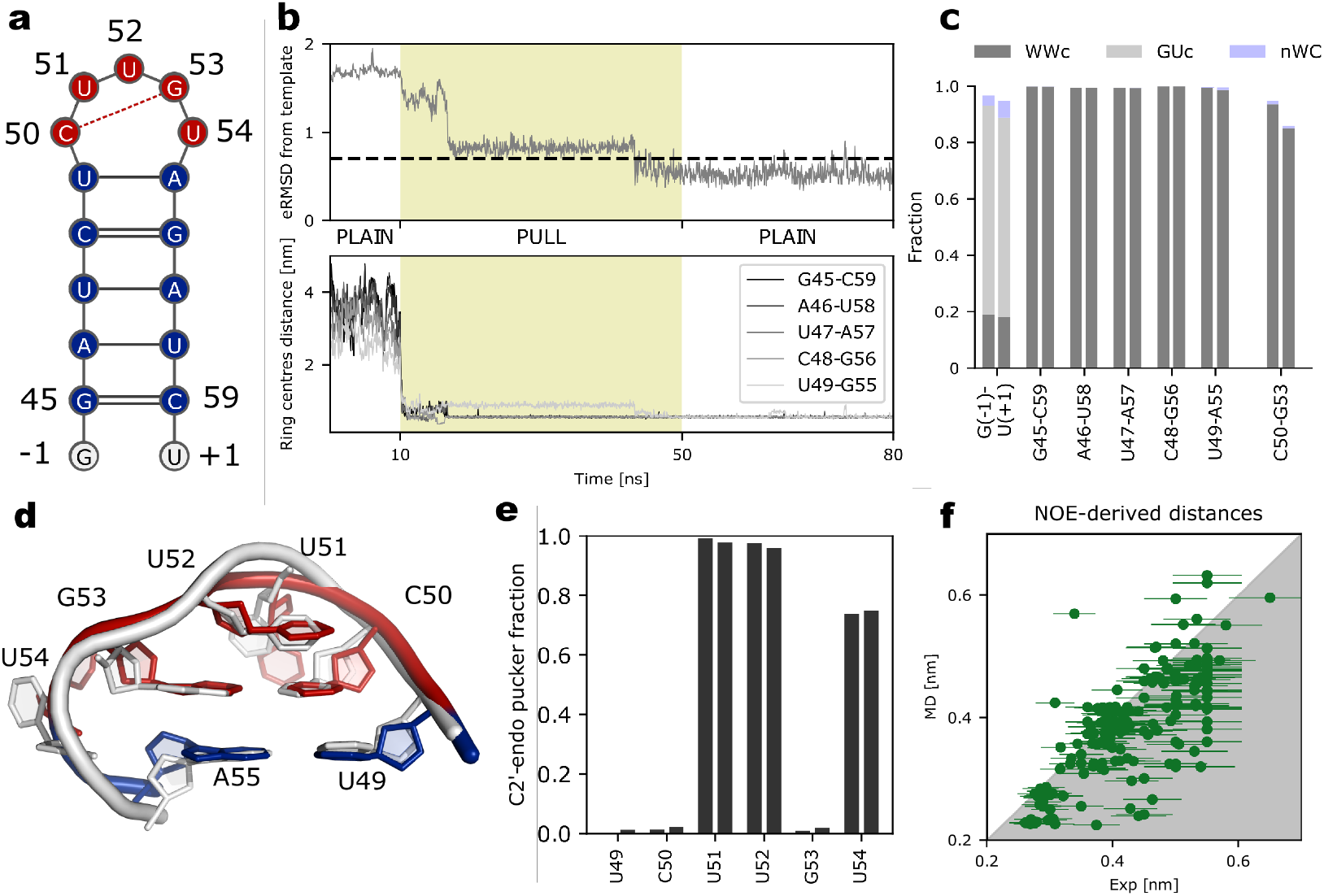
Description of the computational procedure and validation on SL2. **a** Sequence and secondary structure of SL2. The genomic numbering follows the convention in [11]. The region used for constructing the template A-form three-dimensional structure is shown in blue. The sampling in the region shown in red is enhanced by partial tempering. The Watson-Crick base-pair between C50 and G53 predicted in partial tempering simulations is shown as a dashed red line. **b** Behaviour of a successful pulling simulation. eRMSD (top panel) and distance between ring centres of the five WC base-pairs (bottom panel) during the three stages of the procedure: initial plain MD, pulling stage, and final plain MD stage, as labelled. The eRMSD threshold of 0.75 is shown as a dashed horizontal line. **c** Population of base-pairs from 20 replicas × 2.6 *μ*s partial tempering simulations. Following the Leontis-Westhof classification [46], base-pairs are annotated as cis Watson-Crick/Watson-Crick (WWc), GU wobble cis (GUc) or non-Watson-Crick (nWC). For each interaction, the two bars show the statistics from two independent simulations. **d** Centroid of the most-populated cluster from MD simulations (blue/red) superposed onto the SL2 loop from SARS-CoV-1 structure (white) [16]. **e** Sugar pucker population in the loop region. **f** Calculated and experimental NOE-derived upper bound distances. The 10 of the 179 calculated NOEs that are larger than the experimental values are located in the white upper-triangular region.

While pulling simulations make it possible quickly to generate conformations with a desired stem structure, these runs are not sufficiently long to allow rearrangements in the loop region. Previous simulation studies have shown these motions to occur on long timescales (*μ*s to *m*s) that require extensive simulations [47]. We thus enhance the sampling in the loop region (red in Fig. 2a) by running extensive partial tempering simulations (20 replicas, each 2.6 *μ*s-long) initialised from a successful pulling run (see Methods).

Analyses of the enhanced simulations show that all five base-pairs in the stem are stable, and a comparison of two independent simulations shows that our simulations are well converged (Fig. 2c). In addition, we observe the formation of a stable terminal GU wobble base-pair, as well as of a C50-G53 Watson-Crick interaction in the loop, resulting in a structure resembling a CUUG tetraloop [48, 49] with a bulged U at position 54. The interaction between C50 and G53 is also compatible with the decreased sensitive to chemical probing of these two bases relative to the remaining three in the loop [10]. A three-dimensional cluster analysis [50] reveals that the largest cluster (≈ 90%) is remarkably similar to the available NMR loop structure from SARS-CoV-1 [16], whose sequence differs from our construct in the two terminal base-pairs. For example, the centroid of the most populated cluster from MD simulations superimposes extremely well to the first NMR model in PDB 2L6I (Fig. 2d). In the loop, C50 forms a canonical base-pair with G53, U52 stacks on top of C50, and both U51/U54 are solvent exposed. The median distance between the experimental structure and the MD simulation is 1.6 Å heavy-atom root mean square deviation and 0.5 eRMSD, as shown in Supporting Information 1 (SI1).

The sugar puckers of U51, U52, and U54 are preferably in C2’-endo conformation, in full agreement with NMR measurements [16] (Fig. 2e). The great majority (169 out of 179) of upper-bound NOE-derived distances are satisfied in MD simulations (Fig. 2f). The most significant discrepancies consist of short proton-proton distances between G53-U54 that are not sampled in our MD runs. The full list of violated NOEs is shown in Fig. SI2.

### Stem loop 5a

SL5a consists of two helices (blue nucleotides in Fig. 3a) connected by an internal loop and capped by an hexaloop with sequence UUUCGU. We use the same procedure as described above for SL2 to generate initial configurations and to sample the ensemble of SL5a by enhancing regions where secondary structure is not known (red regions in Fig. 3a). Helix 1 is highly stable throughout the 20 replicas × 3.6 *μ*s partial tempering simulations (Fig. 3b).

**Figure 3.**
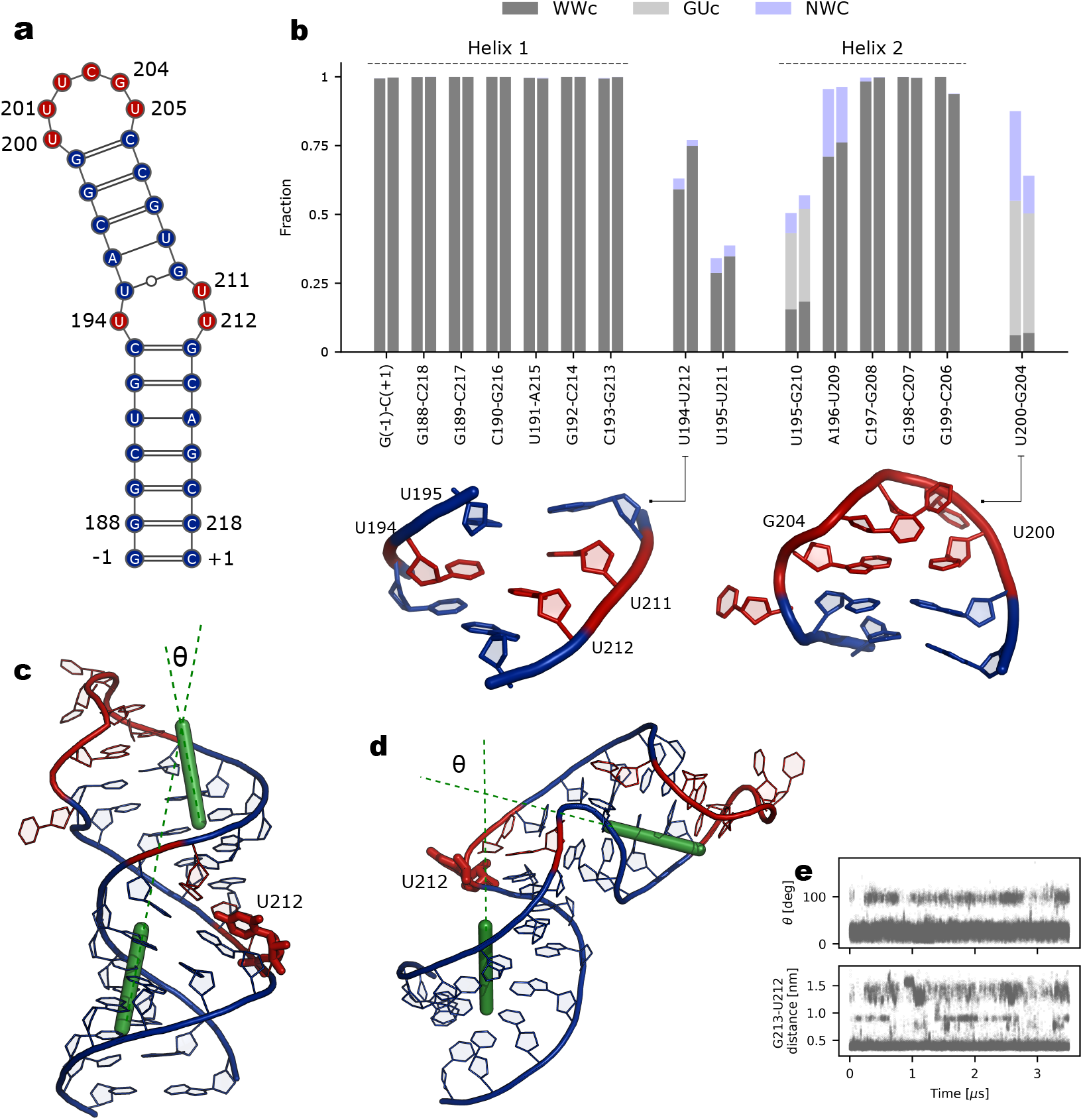
SL5a results. **a** Sequence and secondary structure of SL5a. The genomic numbering follows the convention in [11]. The region used for constructing the A-form templates is shown in blue. The sampling in the region shown in red is enhanced by partial tempering. **b** Population of base-pairs from 20 replicas × 3.6 *μ*s partial tempering simulations. Centroids of the most populated internal and apical loop structures discussed in the text are shown. **c** Representative straight conformation. The axes defined by helix 1 and 2, together with the angle *θ* between them are shown in green. The helical axes are calculated as the vector normal to the base-pair plane, averaged over all base-pairs within the helix. **d** Representative kinked conformation. **e** Time series of the angle between helices, *θ*, (top panel) and of the distance between U212 and G213 (bottom panel) in the neutral replica. We note that due to the replica-exchange setup used in our simulations, these simulations do not reflect the timescales needed to switch between the two conformations.

In the internal loop in SL5a we observe the formation of a U194-U212 base-pair with a population of ≈ 60% (Fig. 3b). The wobble GU base-pair between U195 and G210 in helix 2 is in equilibrium with an alternative WC/WC base-pair between U195 and U211. U212 is highly dynamic (see below), while other nucleotides in the internal loop, including G210, remain in the stack throughout MD simulations. Our predictions are overall compatible with NMR data, suggesting U194 to be base-paired, and U195-G210 at least partially stable [32].

The most frequent conformation of the apical loop is characterised by a wobble interaction between U200 and G204, with U205 completely solvent exposed. Sugar pucker conformation for nucleotides 201 to 204 are predominantly in C2’-endo conformation, at variance with NMR data that indicate C2’-endo conformations only at positions U202 and C203 [32]. In one of the two simulations we observe the transient formation of a U200-U205 base-pair capped by a canonical UUCG tetraloop fold [51]. The low population of this state is not surprising, as the canonical UUCG tetraloop fold has been shown not to be stabilised to a sufficient degree within this force-field [52]. While the detailed architecture of the hexaloop has not yet been solved experimentally [11, 32], our computational prediction and available NMR data cannot be easily reconciled in this case. Instead, we suggest that the simulations might be integrated with structural data collected in the future through a reweighting procedure [40].

In addition to the analysis of individual base-pairs, we identify in the simulations frequent transitions from straight to kinked conformations (Figs. 3c-d). The latter structure is reminiscent of a kink-turn motif, but lacks its signature interactions and sequence propensities [53]. This large-scale motion does not affect the structures of the individual stems, but only their relative orientation. We also note that kinked conformations are observed only when U212 is solvent exposed, and therefore not engaged in base-pairing interactions with U194 or U195 (Fig. 3e), and suggest that the relative populations could be modulated by binding of proteins or small molecules to this motif.

### Stem loop 1

Similarly to SL5a, SL1 consists of two helices connected by an asymmetrical internal loop (Fig. 4a). We use the pulling procedure to initialise our runs from the previously determined secondary structure [11]. While U13-A26/U17-A22 base pairs could not be confirmed by NMR, we have included these two interactions for generating the initial structures as they were present in previous consensus secondary structure assignments [14]. Note that secondary structure restraints are absent during partial tempering simulations, and we indeed observe a rich dynamics involving the loop regions as well as the neighboring residues.

**Figure 4.**
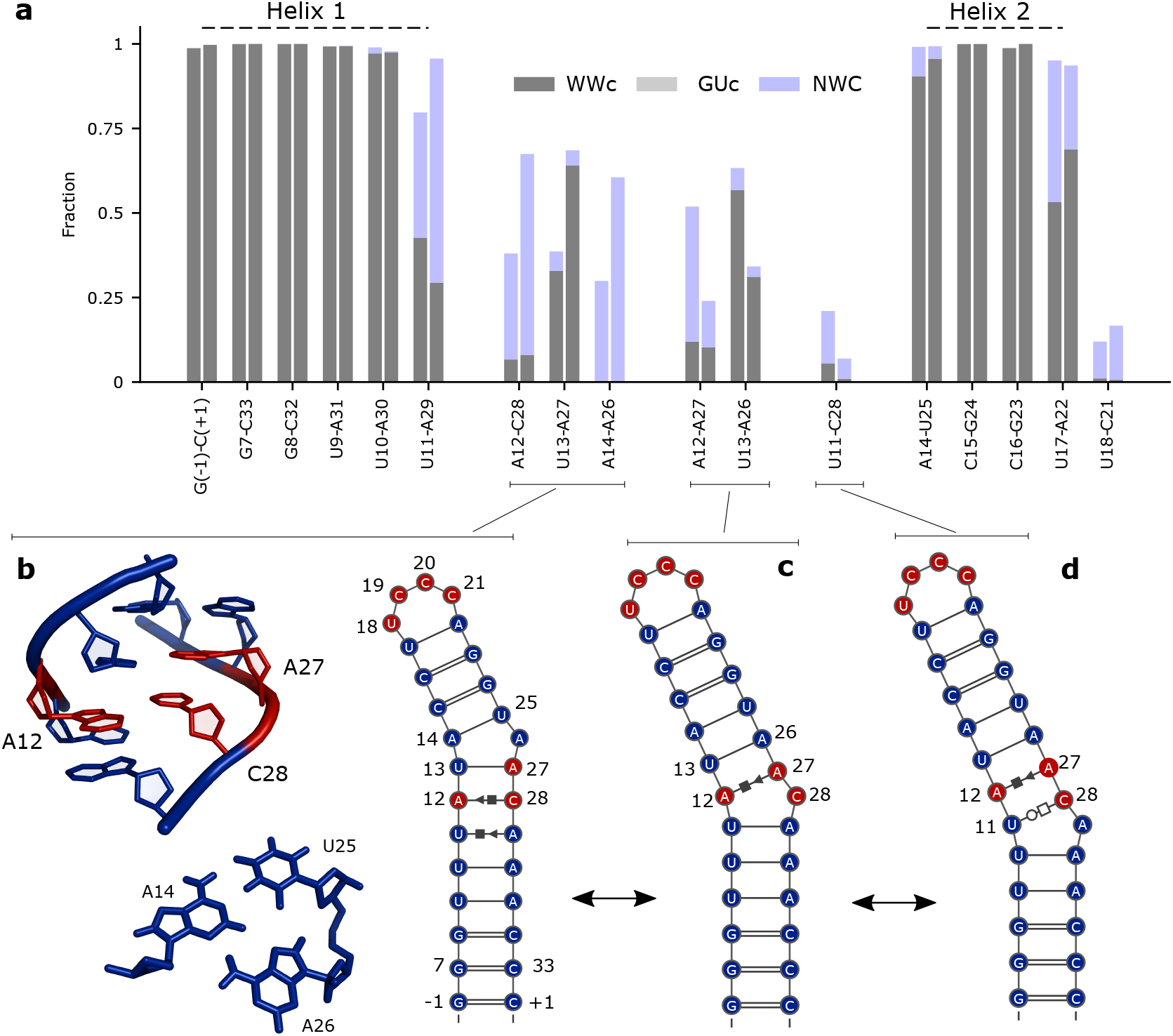
SL1 results. **a** Population of base-pairs from partial tempering simulations. **b** Three-dimensional structure of the most populated internal loop configuration showing the U11-A29 Hogsteen-Sugar, A12-C28 Sugar-Hoogsteen base-pairs, as well as the base A14-U25-U26 base triple. The corresponding secondary structure is shown on the right. The genomic numbering follows the convention in [11]. The base-paired regions selected for creating initial configurations is shown in blue, while partial tempering acts in regions shown in red. **c-d** Secondary structures of two alternative internal loop arrangements.

While helix 1 is stable, the terminal base-pair U11-A29 either interacts through the Watson-Crick/Watson-Crick or Hoogsteen-sugar edges (Fig. 4b). The presence of this non-canonical interaction allows the formation of a A12-C28 sugar-Hoogsteen interaction as well as of a U13-A27 base-pair. A26 is not completely solvent-exposed, but forms a base triple together with A14 and U25. In our simulations we also observe two alternative, lowly populated secondary structures (Fig. 4c-d) where A12-A27/U13-A26 are base-paired, and either C28 or A29 are partially excluded from the stack. Qualitatively, the nucleobase arrangements shown in Fig. 4b-d all preserve the coaxial stacking of the two helices.

The apical loop is highly dynamic and lacks a major, stable conformation with specific base-base interactions. U17-A22 base-pair interconverts from Watson-Crick/Watson-Crick to sugar/Hoogsteen, and U18 can engage a sugar-Hogsteen interaction with C21, albeit with a small population (Fig SI3).

### Stem loop 4

The SL4 element has a more complex organisation, and consists of four A-form helices of different lengths linked by short pyrimidine internal loops and capped by a UGCAU pentaloop (Fig. 5a). The experimentally-derived secondary structure is generally preserved in the parallel tempering simulations (Fig. 5b). The A-form geometry between helix 1 and helix 2 is stabilised by a WC/WC C92-C119 base-pair (Fig. 5b). U95 is solvent-exposed and transiently engages in non-canonical interactions with G94 (Fig. 5b).

**Figure 5.**
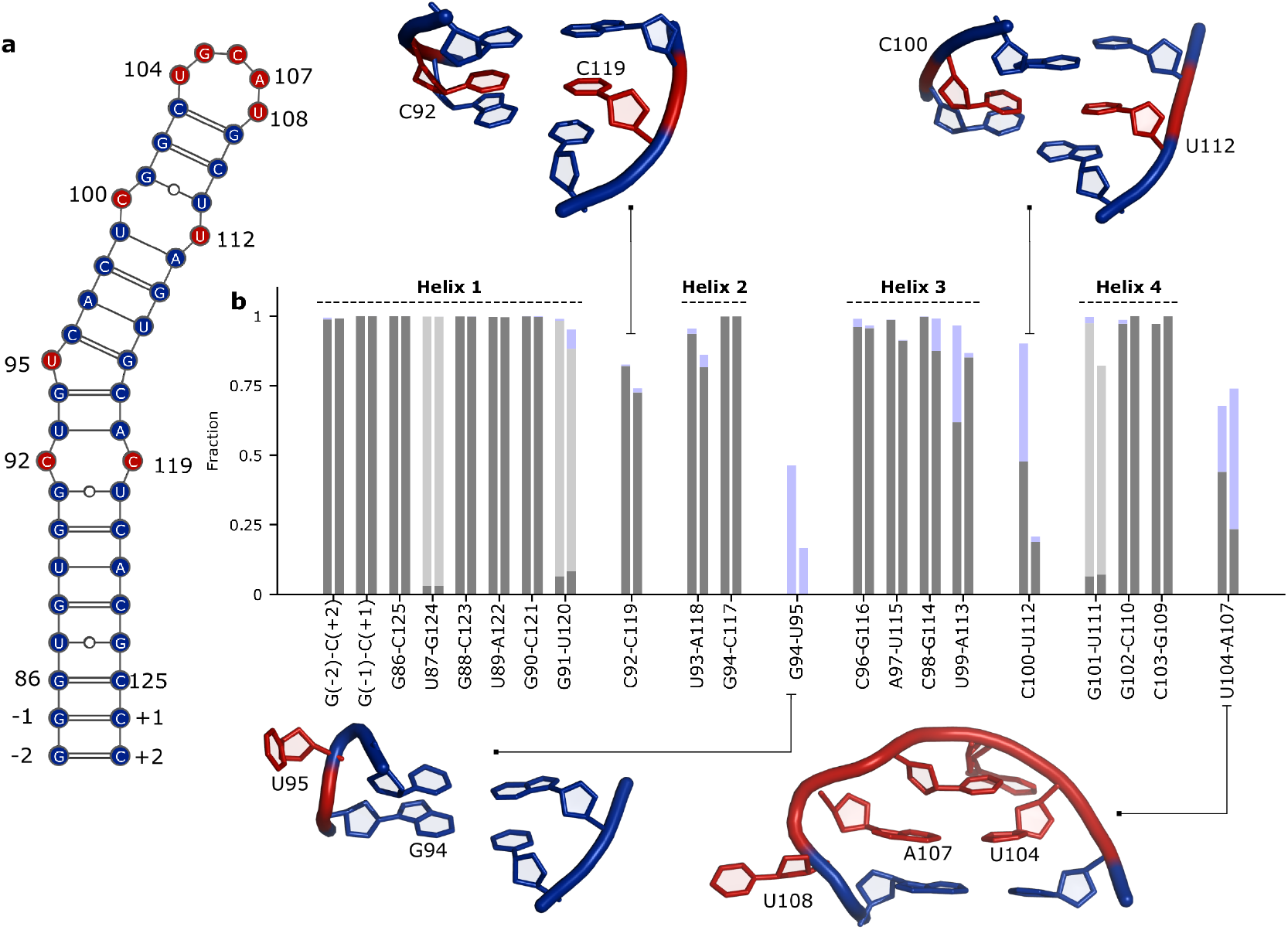
SL4 results. **a** Sequence and secondary structure of SL4. The genomic numbering follows the convention in [11]. The region used for constructing the A-form templates is shown in blue. The sampling in the region shown in red is enhanced by partial tempering. **b** Population of base-pairs from 20 replicas × 1.6 ***μ***s partial tempering simulations. Centroid of the most populated internal and apical loops discussed in the text are shown.

While the interactions observed in the first and second loops are relatively consistent between the two independent simulations, this is not the case for the the third internal loop. Here, the two simulations behave differently suggesting slow motions that are not fully sampled in our simulations. In the first run C100 preferentially base-pairs with U112, while in the second run additional configurations are sampled, where either C100 or U112 are not in the stack. The partial overlap between the two runs, that we ascribe to insufficient sampling, is further confirmed by a principal component analysis of this internal loop region (SI4), as well as from the low number of round trips in temperature space in run 1 (SI5). Experimental NMR measurements on SL4 agree with A-form helix compatible arrangement of the opposing residues C100 and U112 [11], thus supporting the presence of pairing between the two bases. The apical loop structure resembles a tetraloop with U108 solvent-exposed and U104:A107 forming either a Watson-Crick/Watson-Crick or a sugar/Hogsteen non-canonical interaction, as shown in Fig. 5b.

### SL2+SL3

We here report the results on the SL2+SL3 construct, consisting of two consecutive stem-loop structures: SL2 (described above) and SL3, which contains the important transcription-regulating sequence required for subgenomic viral RNA synthesis [37]. Although we did not enforce this geometry, we find structures where the two helices are stacked coaxially tail-to-tail, thereby forming a unique A-form-like stem (Fig. 6).

**Figure 6.**
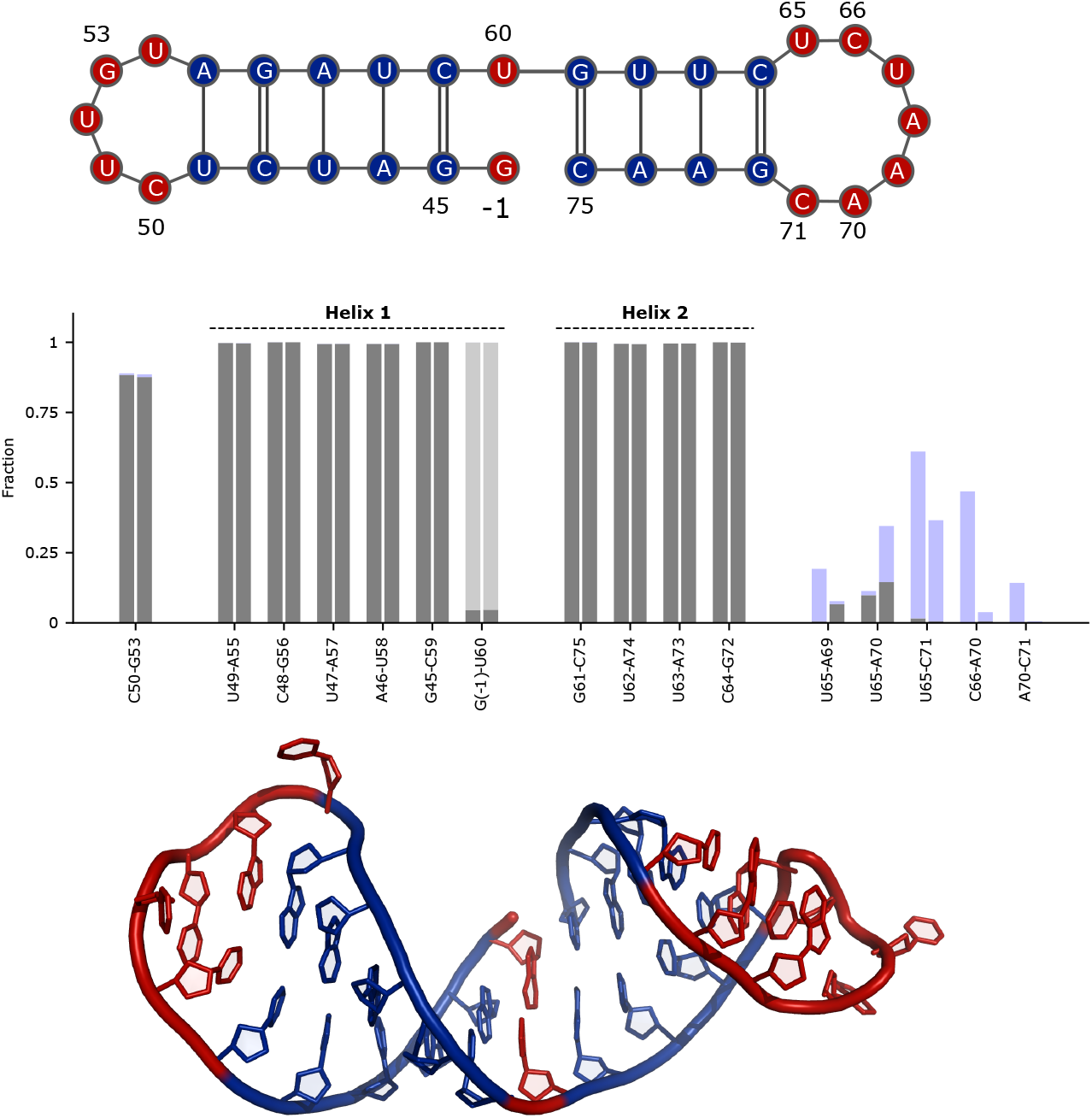
SL2+SL3 results. Top: Sequence and secondary structure of SL2+SL3. The genomic numbering follows the convention in [11]. The region used for constructing the A-form templates is shown in blue. The sampling in the region shown in red is enhanced by partial tempering. Middle panel: Population of base-pairs from partial tempering simulations. Bottom panel: Centroid of the most populated cluster.

The SL2 loop conformations are similar to those observed for SL2 alone, with a stable C50-G53 base-pair (see above). The UCUAAAC heptaloop in SL3 is highly flexible, and adopts several conformations that are partially structured in terms of base-pairing (Fig. 6). A cluster analysis of SL3 conformations reveals the presence of a loop structure where U65 forms a stacking interaction with C66, followed by the strand inversion and by two consecutive stackings between A68-A69-A70. This specific loop architecture can be considered the most RNA-like sampled in simulations, as we found it to be remarkably similar to Helix 58 loop in the archeal large ribosomal subunit [54] as well as to the anticodon loop in a nonproteinogenic tRNA [55] (Fig. SI6).

Helix 2, containing the transcription regulatory sequence ACGAAC is fully stable in our simulations. NMR measurements show that helix 2 is less stable than helix 1 in the absence of magnesium [11]. Such decreased stability of helix 2 is in line with a model for the body-to-leader fusion mechanism, in which the leader transcription regulatory sequence, ‘hidden’ inside SL3, is required to unfold to be active [3]. The tail-to-tail structure observed in simulations, which may not form in the context of the full 5’-UTR construct, may overstabilise helix 2 in this construct. Note that quantifying duplex stability is a challenging problem that is not considered here, as it usually requires dedicated simulation protocols [47].

## Conclusions

We report the prediction of the structure and conformational heterogeneity of five stem-loop elements, ranging in size between 17 and 44 nucleotides, located at the 5’-UTR in SARS-CoV-2 genome. As a starting point for our study we use the secondary structure determined by NMR experiments [11].

Structures of several of these elements have also been predicted using FARFAR2 [17]. A principal component analysis of the structures generated for SL1 and SL2 shows small overlap between FARFAR2 and MD ensembles, suggesting the two methods to be complementary (SI7). In the case of SL2, the MD predictions mostly agree with the available NMR data, albeit small discrepancies remain. For this structure FARFAR2 does not predict the experimental loop architecture with the C50-G53 base-pair (SI1), and it thus appears that the ensemble generated by MD provides a more accurate description in this case. Given the scarcity of available experimental data on the 5’-UTR elements it is not possible at this stage to compare the accuracy of different computational methods. Indeed, FARFAR2 has been shown to better predict experimental data compared to MD on an RNA internal loop [28].

As in most MD studies, the results presented in this work depend on the amount of sampling. We took care in identifying and discussing states that are visited multiple times during one simulation and in both replicates, indicating these conformations to correspond to *bona fide*, highly populated metastable states. Moreover, the overall high agreement between the two independent simulations suggests that the conformational ensembles are relatively well converged. Our simulations of SL4, consisting of 44 nucleotides, represents an important exception with greater deviations between the two simulations, in particular of the third internal loop (SI4, SI5), indicating that our approach would require larger computational resources for molecules of this size.

The modelling protocol used in the present study implicitly considers helix formation and internal loop dynamics as partially decoupled events. While this is a relatively common assumption in computational studies of RNA [18, 31, 27], we can envisage situations in which the rearrangement of a loop requires one or more base pairs to break transiently. In our partial tempering MD simulations the secondary structure is not fixed: these rare events can thus be observed, albeit with a low frequency due to the absence of auxiliary replicas enhancing the breakage and formation of stem regions.

Despite recent improvements of RNA force-fields [56, 47, 43], the accuracy of the simulation results still depend on the accuracy of the physical models used, and we have validated our simulations against available experimental data. As more data becomes available, we note that it should be straightforward to integrate our extensive conformational sampling with data from future solution measurements (e.g. NMR or SAXS) [38, 39, 40] to refine the atomic-level description of SARS-CoV-2 RNA structure and dynamics.

## Methods

MD simulations were conducted using the Amber ff99SB force field with parmbsc0 [57] and OL3 [58] corrections in conjunction with OPC water model [59]. The initial conformations were minimised in vacuum and then solvated in a dodecahedral box with minimum distance from box edge of 1.2nm. Potassium ions were added to neutralise the box [60]. The systems were equilibrated in the NVT and NPT ensembles at 1 bar for 1 ns. Production runs were performed in the canonical ensemble using a stochastic velocity rescaling thermostat [61]. All bonds were constrained using the LINCS [62] algorithm and equations of motion were integrated with a time step of 2 fs. Simulations were performed using GROMACS 2019.4 [63] patched with PLUMED 2.5.3 [64]. Force-field parameters are available at http://github.com/srnas/ff/tree/opc-water-model/amber_na.ff.

### Pulling simulations

The aim of the pulling simulations is quickly to generate three-dimensional RNA conformations with a specific input secondary structure. We adapt a procedure originally introduced for reconstructing atomistic structures from coarse-grained models [25]. Our protocol consists of three stages, as illustrated in Fig. 2a-b for SL2:

1. Plain MD simulation initialised from extended, single stranded, A-form RNA structures.
2. Pulling. We construct a reference structure with an ideal A-form double stranded RNA template for regions with known secondary structure (shown in blue in Fig. 2a) and perform pulling simulations to minimise the eRMSD [45] distance between the RNA molecule and the reference. We here use adiabatic bias MD simulations [65], that introduce a biasing potential damping the fluctuations only when the system moves further away from the template. In this way the pulling procedure follows the thermal fluctuations of the system, and does not require a pre-defined scheduling for the moving restraint [25]. When more than one stem is present, the bias acts on multiple templates simultaneously. The constant of the harmonic damping bias is set to 2 kJ/mol.
3. Plain MD. After 40ns of pulling simulation, the external bias is removed and the system freely fluctuates without external forces.

At the end of the procedure we extract samples from the final plain MD stage satisfying the secondary structure constraints, i.e. with eRMSD from template < 0.75. This criterion guarantees that all base-pairs corresponding to those in the target structure are correctly formed (Fig. 2b, bottom panel). Note that individual pulling simulations may fail: stems are sometimes not formed correctly and unfold during the final plain MD stage (Fig. SI8). The success rate of this procedure ranges from ≈ 80% for SL2 to ≈ 10% for SL4, which consists of 4 stem structures, consistent with previously reported benchmarks [25]. During preliminary tests we have observed that the formation of the correct secondary structure is hampered by early-forming non-native contacts. For this reason we conduct pulling simulations at 340K.

### Partial tempering

We resolvate and minimise the RNA structures obtained from pulling as described above. We use partial tempering to enhance sampling in regions with unknown secondary structure [66, 26]. One reference replica exchanges via the Metropolis-Hastings algorithm with parallel simulations that are conducted with a modified Hamiltonian. The modification is such that interactions in the cold region (blue and grey in Fig. 2a, together with ions and water molecules) are kept at the reference temperature T=300K. Interactions in the hot region (shown in red in Fig. 2a) are rescaled by a factor 1/*λ*, while the interactions between cold and hot regions are at an intermediate effective temperature of 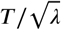, with 1 ≤ *λ* < 0. In this study we use 20 geometrically distributed replicas in the effective temperature range of 300 to 1000 K, and attempting exchanges every 240 steps. Hot regions are shown in red in Figs. 2–6, and include the phosphate group at the 3’-end of the selected region. We have found this small addition to be beneficial in preliminary test runs. System length, simulation time per replica, average acceptance rate, and number of round trips in temperature space are reported in Table 1, and additional statistics are available in Fig. SI5. To help the identification of slow degrees of freedom and assess convergence we conduct two independent runs with different initial conformations obtained from two separate pulling simulations (Fig. SI9).

**Table 1.**
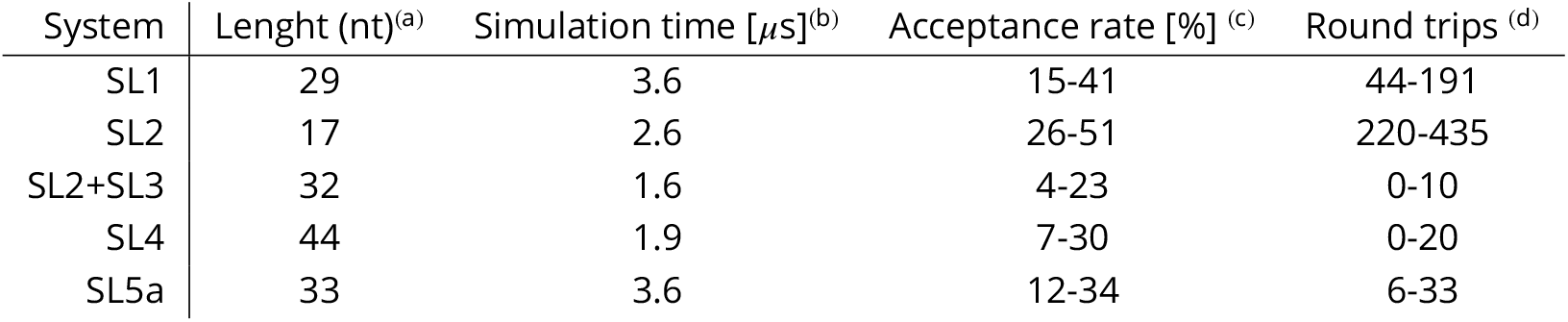
Simulated systems and related statistics. (a) number of nucleotides (nt). (b) Per-replica simulation time. Each simulation consists of 20 replicas and two independent runs for each system are carried out. The first 100ns of each run are discarded during analysis. (c) Minimum and maximum acceptance rate across all replicas and runs (d) Minimum and maximum number of round trips in temperature space across all replicas and runs.

### Analysis and visualisation

Ideal A-form structures were constructed using the nucleic acids builder (NAB) software in Ambertools [67]. Trajectories were analysed using PLUMED [64] and MDtraj[68]. Base-pair annotation and puckering angles were calculated with Barnaba [50].

## Supporting Information

1. Root mean squared deviation between PDB:2L6I, MD and FARFAR2 simulations.
2. NOE measurements violated in partial tempering simulations.
3. Three-dimensional centroids of the first five clusters in SL1 apical loop.
4. Partial tempering simulations of SL4 projected onto the first two principal components.
5. Average acceptance rate and number of round trips in temperature space in partial tempering simulations.
6. SL3 structure similar to experimental loop structures in the PDB database.
7. Principal component analysis performed on FARFAR2 and MD ensembles for SL1 and SL2.
8. Example of unsuccessful pulling simulation.
9. Initial configurations in partial tempering simulations.

## Acknowledgements

We acknowledge Drs. Harald Schwalbe, Anna Wacker, Julia E. Weigand, Andreas Schlundt, and other members of the Covid19-NMR consortium for discussions and insights. SB and KLL acknowledge funding from the Lundbeck Foundation BRAINSTRUC structural biology initiative (R155-2015-2666). We acknowledge access to computational resources from PRACE for the COVID-RNA project (COVID19-72).

## Data Availability

Analysis scripts, MD trajectories, and input files are available via https://github.com/KULL-Centre/papers/tree/master/2020/COVID_5UTR_MD PLUMED input files are available in the PLUMED NEST [69] under the accession code 20.034 https://www.plumed-nest.org/eggs/20/034/.

