## Supplementary information for "Conformational Ensembles of Non-Coding Elements in the SARS-CoV-2 Genome from Molecular Dynamics Simulations"

(1) Structural Biology and NMR Laboratory & Linderstrøm-Lang Centre for Protein Science Department of Biology, University of Copenhagen. Ole Maaløes Vej 5, DK-2200 Copenhagen N, Denmark
(2) Scuola Internazionale Superiore di Studi Avanzati (SISSA), Via Bonomea 265, 34136, Trieste, Italy

### SUPPORTING INFORMATION

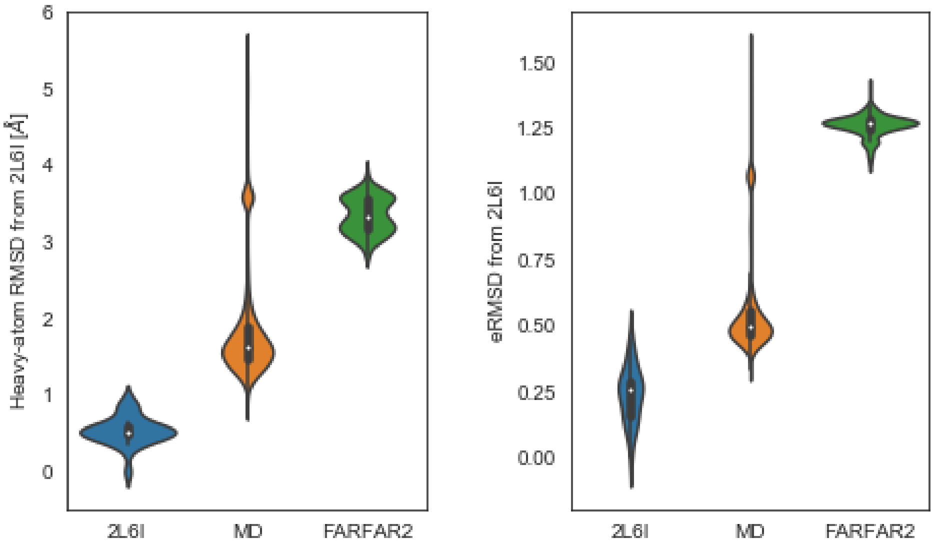


**SI1** Left: Heavy-atom root mean squared deviation distribution between model 1 in PDB:26LI and different ensembles: 2l6I bundle [1], partial tempering MD simulations, and FARFAR2 models [2]. The same distances calculated using the eRMSD [3] are shown in the right panel.

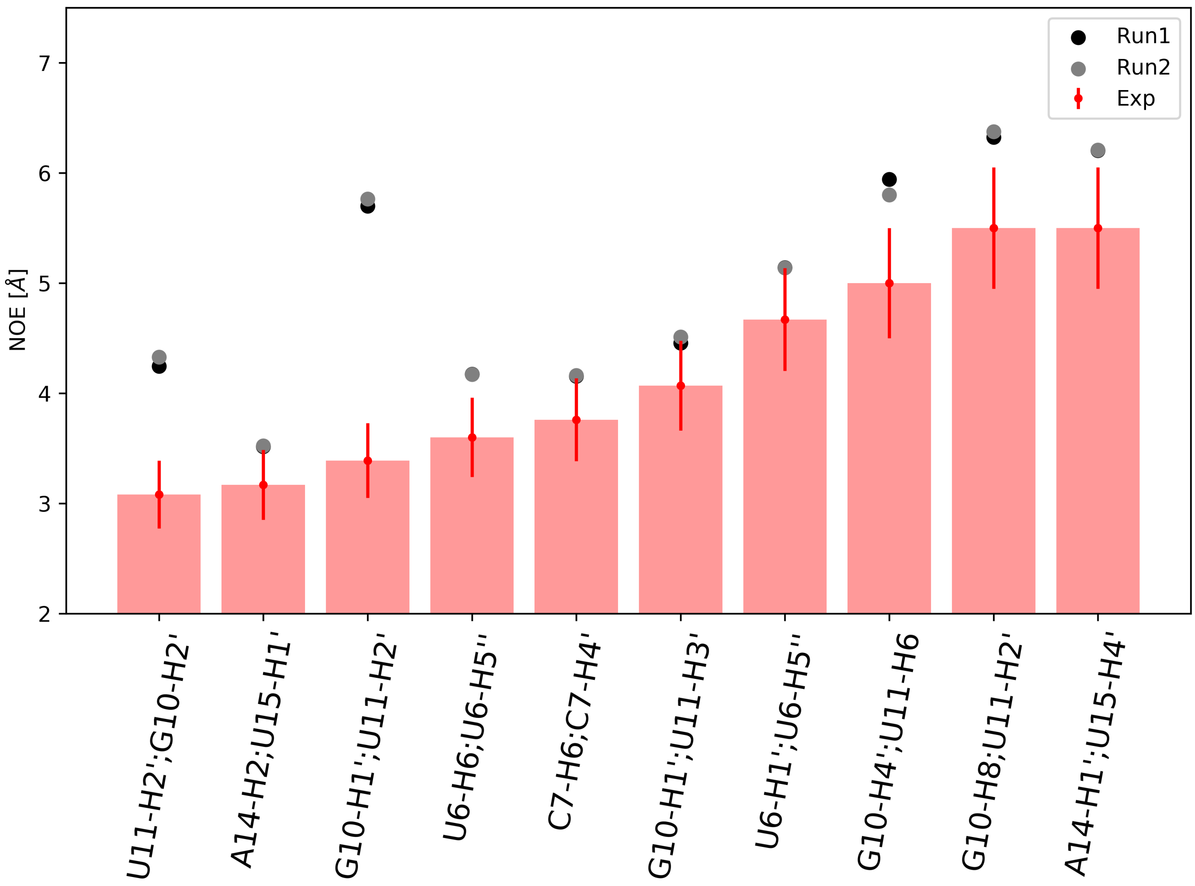


**SI2** Barplot showing the list of the 10 upper-bound NOE measurements violated in partial tempering simulations. Averages from both runs are reported.

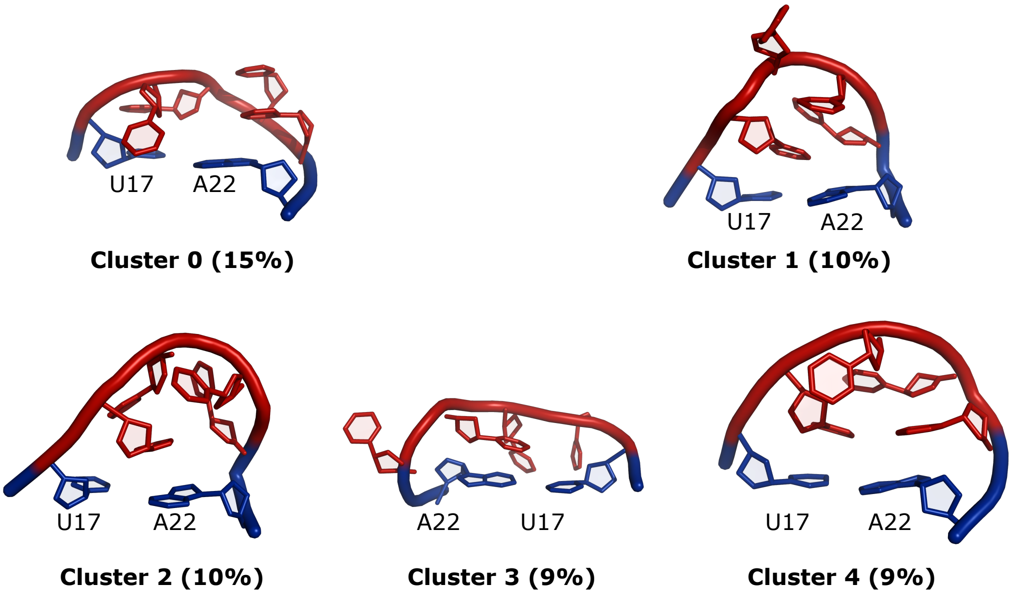


**SI3** Three-dimensional centroids and corresponding populations of the first five clusters in SL1 apical loop.

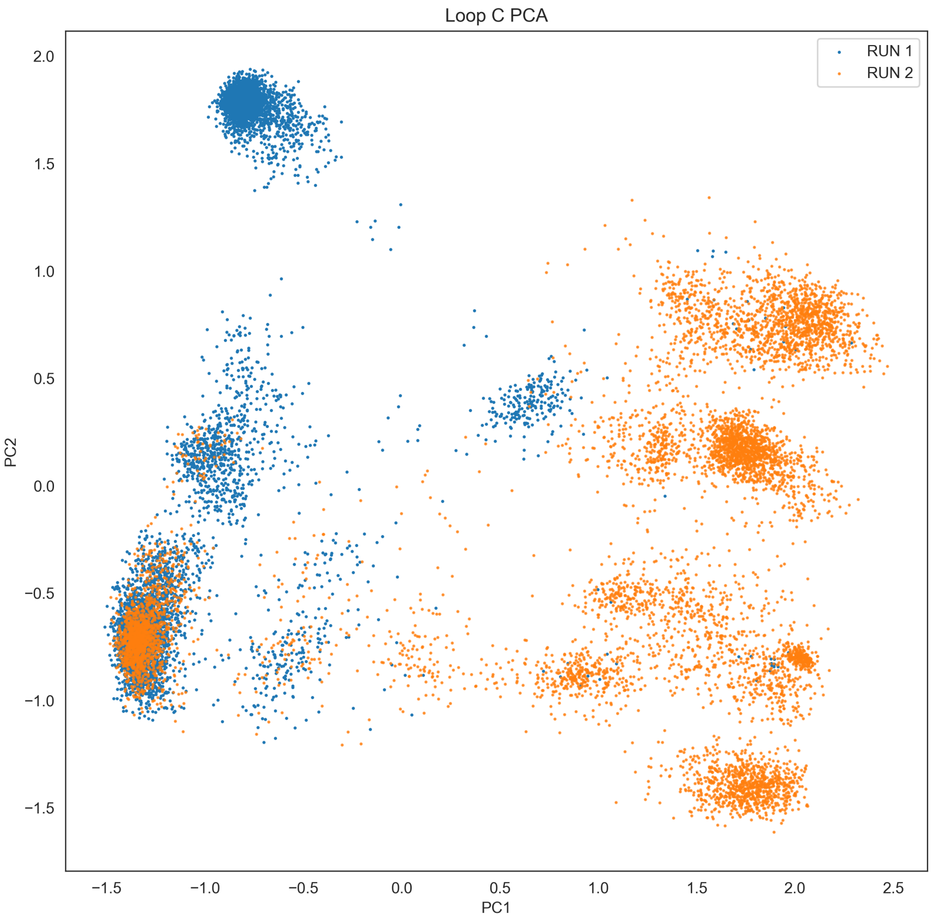


**SI4** Partial tempering simulations of SL4 projected onto the first two principal components calculated as described in [4]. Only residues in the internal loop region (99-100-101-111-112-113) are considered.

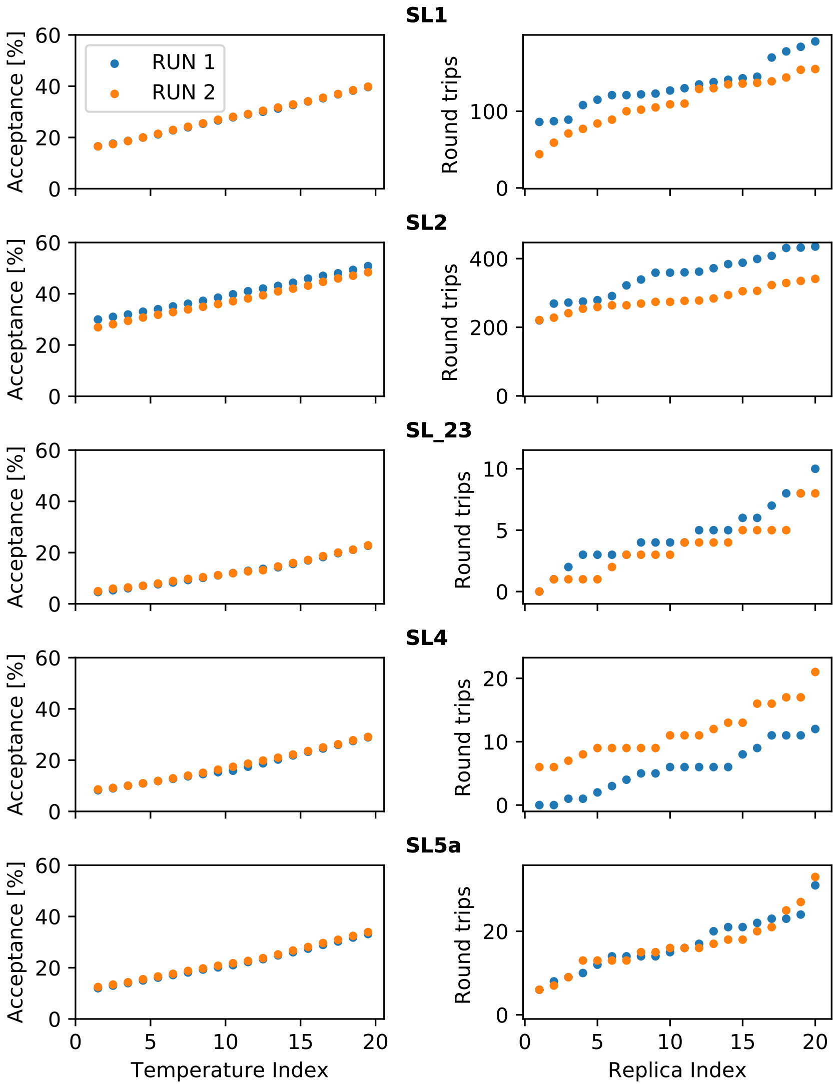


**SI5** Left panels: average acceptance rate between temperatures in partial tempering simulations. Right panels: number of round trip times in temperature space for each replica. A replica completes one round trip when going from temperature 1 to temperature 20 and back. Statistics for each system and for both runs are reported in the five rows, as labeled.

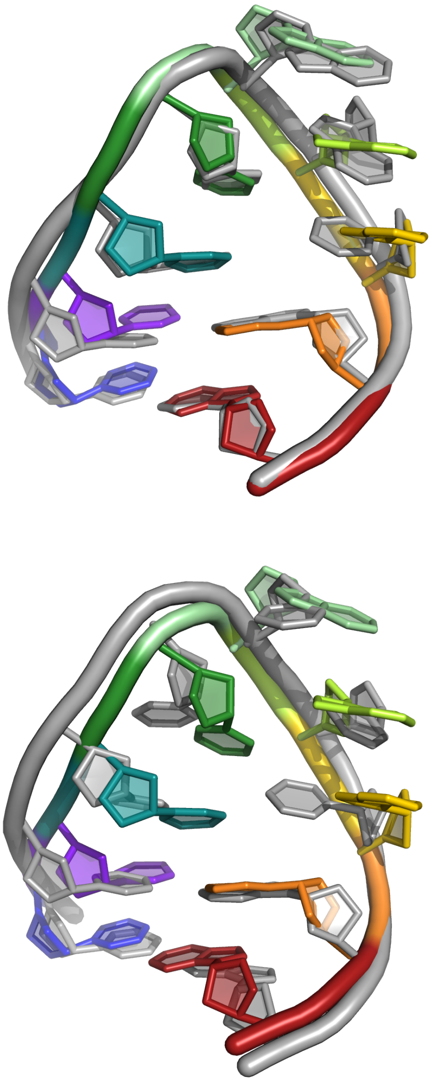

**SI6** Top: Cluster centroid 2 from SL2+SL3 simulation (in color) superposed onto helix 58 loop from H.Marismortui large ribosomal subunit (PDB 1FFK residues 1593 to 1601, in gray). Bottom: Cluster centroid 2 from SL2+SL3 simulation (in color) superposed onto the tRNA anticodon loop (PDB 2LBK residues 5-13, gray).

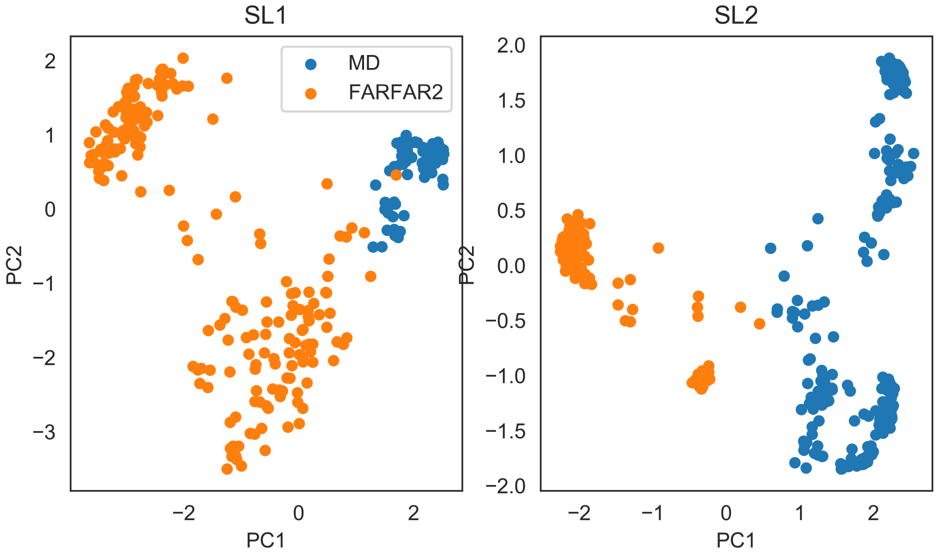


**SI7** Principal component analysis performed on FARFAR2 [2] and MD ensembles for SL1 and SL2. A number of random samples are extracted from the MD ensembles so as to match the number of FARFAR2 models.

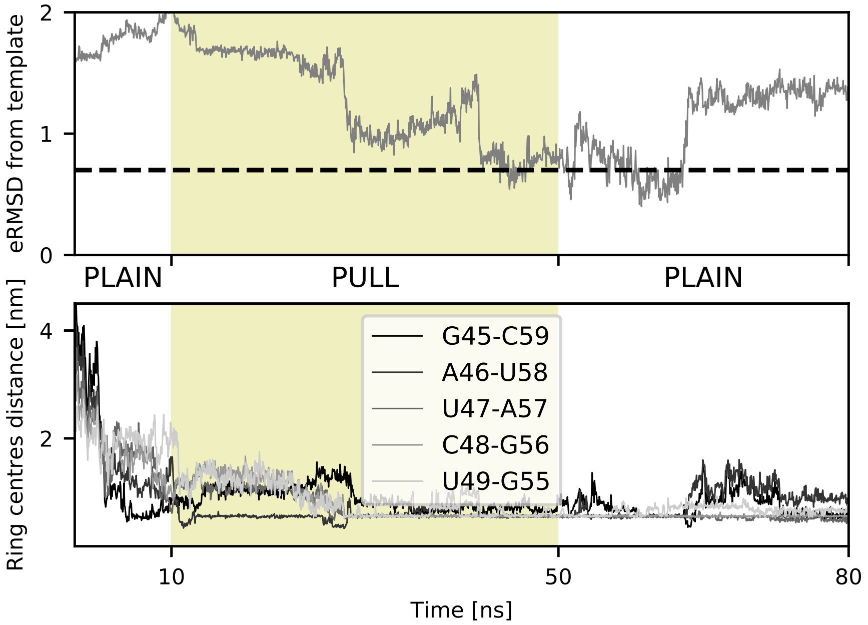


**SI8** Example of an unsuccessful pulling simulation on SL2. While several base-pairs are formed during the pulling stage, partial unfolding occurs during the final plain MD stage.

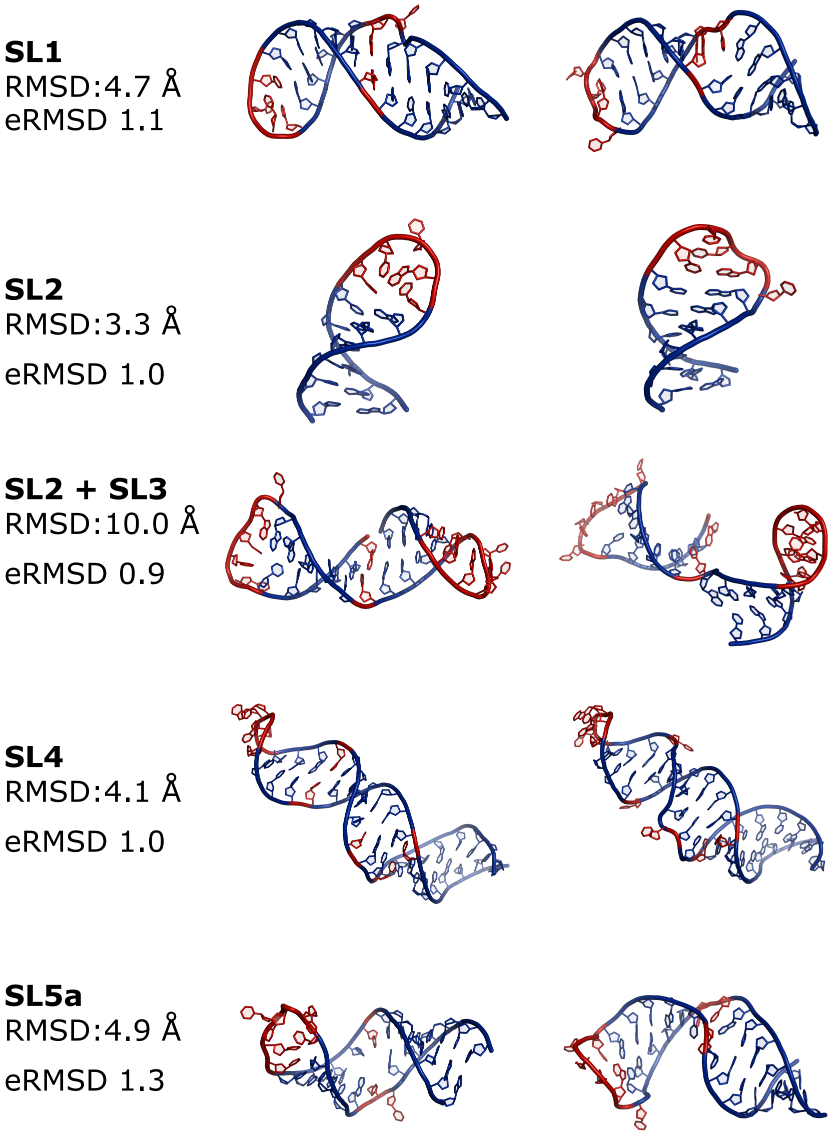


**SI9** Three-dimensional structures obtained from pulling simulations. These structures are used as initial conformations for two independent partial tempering runs. The heavy-atom RMSD and eRMSD between the two are reported.

[2] Ramya Rangan, Andrew M Watkins, Wipapat Kladwang, and Rhiju Das. De novo 3d models of sars-cov-2 rna elements and small-molecule-binding rnas to guide drug discovery. *bioRxiv*, 2020.

[3] Sandro Bottaro, Francesco Di Palma, and Giovanni Bussi. The role of nucleobase interactions in RNA structure and dynamics. *Nucleic Acids Res*, 42(21):13306–13314, 2014.

[4] Sandro Bottaro, Giovanni Bussi, Giovanni Pinamonti, Sabine Reißer, Wouter Boomsma, and Kresten Lindorff-Larsen. Barnaba: software for analysis of nucleic acid structures and trajectories. *Rna*, 25(2):219–231, 2019.
